# MicroRNAs contribute to the host response to *Coxiella burnetii*

**DOI:** 10.1101/2022.05.11.491587

**Authors:** Madhur Sachan, Katelynn R. Brann, Daniel E. Voth, Rahul Raghavan

## Abstract

MicroRNAs (miRNAs), a class of small non-coding RNAs, are critical to gene regulation in eukaryotes. They are involved in modulating a variety of physiological processes, including the host response to intracellular infections. Little is known about miRNA functions during infection by *Coxiella burnetii*, the causative agent of human Q fever. This bacterial pathogen establishes a large replicative vacuole within macrophages by manipulating host processes such as apoptosis and autophagy. We investigated miRNA expression in *C. burnetii*-infected macrophages and identified several miRNAs that were down- or up-regulated during infection. We further explored the functions of miR-143-3p, an miRNA whose expression is down-regulated in macrophages infected with *C. burnetii*, and show that increasing the abundance of this miRNA in human cells results in increased apoptosis and reduced autophagy – conditions that are unfavorable to *C. burnetii* intracellular growth. In sum, this study demonstrates that *C. burnetii* infection elicits a robust miRNA-based host response, and because miR-143-3p promotes apoptosis and inhibits autophagy, down-regulation of miR-143-3p expression during *C. burnetii* infection likely benefits the pathogen.

## INTRODUCTION

The highly infectious intracellular pathogen *Coxiella burnetii* is the etiological agent of Q fever (1–3). After uptake by a host cell, typically an alveolar macrophage, *C. burnetii* establishes a replicative niche termed the *Coxiella*-containing vacuole (CCV) that matures by fusing with lysosomal, autophagic, and secretory vesicles (4–6). To facilitate intracellular replication, *C. burnetii* secretes effector proteins into the host cytosol using a Dot/Icm Type IVB secretion system (T4SS) that disrupt several host processes, including apoptosis, lipid metabolism, inflammation, and vesicular trafficking (3, 7–12). For instance, *C. burnetii* inhibits host apoptosis by recruiting anti-apoptotic Bcl-2 to the CCV, inactivating pro-apoptotic Bad, and promoting a pro-survival response by activating Erk1/2, Akt, and PKA signaling (13–16). Similarly, autophagy-related proteins such as LC3 and p62 are recruited to CCV in a T4SS-dependent manner (10, 17), and inhibition of autophagy components results in reduced *C. burnetii* replication (18, 19).

MicroRNAs (miRNAs) are a class of single-stranded, small (∽22 nucleotides), non-coding RNAs that orchestrate post-transcriptional gene regulation in eukaryotes and some viruses (20). In humans, miRNAs regulate a large number of genes, primarily by inhibiting target gene expression via translation repression and messenger RNA (mRNA) degradation (21–23). Studies have shown that miRNAs are integral to the host response to bacterial, viral, and parasitic infections (24–31); however, miRNAs could either promote or inhibit infection. For example, induction of miR-142-3p expression in macrophages leads to down-regulation of N-Wasp, an actin-binding protein, resulting in reduced uptake of *Mycobacterium tuberculosis* (32). Conversely, induction of miR-125a-3p in macrophages inhibits autophagy and phagosomal maturation, which favors intracellular survival of *M. tuberculosis* by promoting autophagosomal escape (33).

Expression of miRNAs is perturbed in macrophages infected with *C. burnetii* (34), but their potential roles are unknown. In this study, we investigated macrophage gene expression across five time-points during *C. burnetii* infection and identified a large number of miRNAs and protein-coding genes that were differentially expressed, suggesting their involvement in the host response to infection. We also demonstrate that transfecting host cells with miR-143-3p results in reduced *C. burnetii* growth, enhanced apoptosis and diminished autophagy. Collectively, our data indicate that an intracellular environment with a low level of miR-143-3p is conducive to *C. burnetii* growth.

## RESULTS

### Host gene expression correlates with bacterial growth

To identify infection-associated miRNAs and their potential mRNA targets, we measured miRNA and protein-coding gene expression in THP-1 macrophages infected with *C. burnetii* Nine Mile RSA439 Phase II (NMII). The total number of differentially expressed (log2 fold-change ≥ 0.75, Padj ≤ 0.05) miRNAs and mRNAs increased from 25 to 60 and 454 to 6,525, respectively, from day 1 to day 3 (**Table 1 and S1**). By day 5, far fewer miRNAs and protein-coding genes ((34, 211, respectively) were up- or down-regulated in *C. burnetii*-infected cells. This pattern of gene expression suggests that the magnitude of the host cell response to *C. burnetii* infection increases as the bacterium actively replicates (day 3), and as LCVs transition to metabolically less active SCVs (day 5) (35), host response becomes muted in tandem.

**Table 1.** Number of differentially-expressed miRNAs and mRNAs in *C. burnetii*-infected THP-1 macrophages.

|  | **miRNA<sup>a</sup>** |  |  | **mRNA<sup>a</sup>** |  |  |
| --- | --- | --- | --- | --- | --- | --- |
| **Time** | **Total<sup>b</sup>** | **Down** | **Up** | **Total<sup>b</sup>** | **Down** | **Up** |
| 8 hpi | 25 | 12 | 13 | 454 | 220 | 234 |
| 24 hpi | 25 | 16 | 9 | 1,160 | 665 | 495 |
| 48 hpi | 35 | 23 | 12 | 1,742 | 445 | 1,297 |
| 72 hpi | 60 | 43 | 17 | 6,525 | 3,236 | 3,289 |
| 120 hpi | 34 | 18 | 16 | 211 | 80 | 131 |

<sup>a</sup>The current build of the human genome (GRCh38) has 2,654 mature miRNA sequences and 19,231 protein-coding genes.
<sup>b</sup>Differential expression ( $\log_2$ fold change $\geq 0.75$ ; $P_{adj} \leq 0.05$ , Wald test) at respective hours post-infection (hpi) was calculated using DESeq2 (72).

### miRNAs potentially regulate multiple host signaling pathways during *C. burnetii* infection

To identify genes and metabolic pathways targeted by miRNAs, we first performed inverse-expression pairing using the Ingenuity Pathway Analysis (IPA) tool (36). We selected miRNAs and their known or predicted target genes that showed an inverse pattern of expression in *C. burnetii*-infected cells. For instance, if expression of an miRNA is up-regulated in *C. burnetii*-infected cells, we chose targets that are down-regulated, and vice versa. This analysis identified 14, 18, 25, 51, and 23 miRNAs at 8 h, 24 h, 48 h, 72 h, and 120 h post-infection (hpi), respectively, that were inversely paired with differentially-expressed target mRNAs **(Table S2)**. We then investigated biochemical pathways that were enriched for these proteins using the Core Analysis function in IPA, which revealed 215 pathways, including apoptosis signaling, PI3K/AKT, and autophagy that are likely regulated by miRNAs **(Figure 1, Table S3)**. To investigate the role of miRNAs in apoptosis, a process known to be important during *C. burnetii* infection (13, 37–40), we measured expression of 84 miRNAs associated with apoptosis using a targeted qPCR array (41). This assay showed that 12 miRNAs were up- or down-regulated in *C. burnetii*-infected cells at 72 hpi (**Table 2**), suggesting their involvement in apoptosis regulation as part of the host response to *C. burnetii*.

**Table 2.** Differentially-expressed (p ≤ 0.05, n = 3) miRNAs in NMII-infected THP-1 cells compared to uninfected controls.

| miRNA | Fold Change | p-value |
| --- | --- | --- |
| hsa-miR-708-5p | 0.6225 | 0.037471 |
| hsa-miR-145-5p | 0.6312 | 0.006722 |
| hsa-miR-143-3p | 0.6535 | 0.007199 |
| hsa-miR-106b-5p | 0.7386 | 0.032799 |
| hsa-miR-181d-5p | 0.7611 | 0.028265 |
| hsa-miR-16-5p | 0.8026 | 0.024211 |
| hsa-miR-222-3p | 0.8804 | 0.014724 |
| hsa-miR-365b-3p | 1.1015 | 0.016939 |
| hsa-miR-218-5p | 1.5832 | 0.013546 |
| hsa-miR-125a-5p | 1.5868 | 0.010438 |
| hsa-miR-192-5p | 1.7689 | 0.03232 |
| hsa-miR-146a-5p | 5.9635 | 0.000006 |

**Figure 1.**
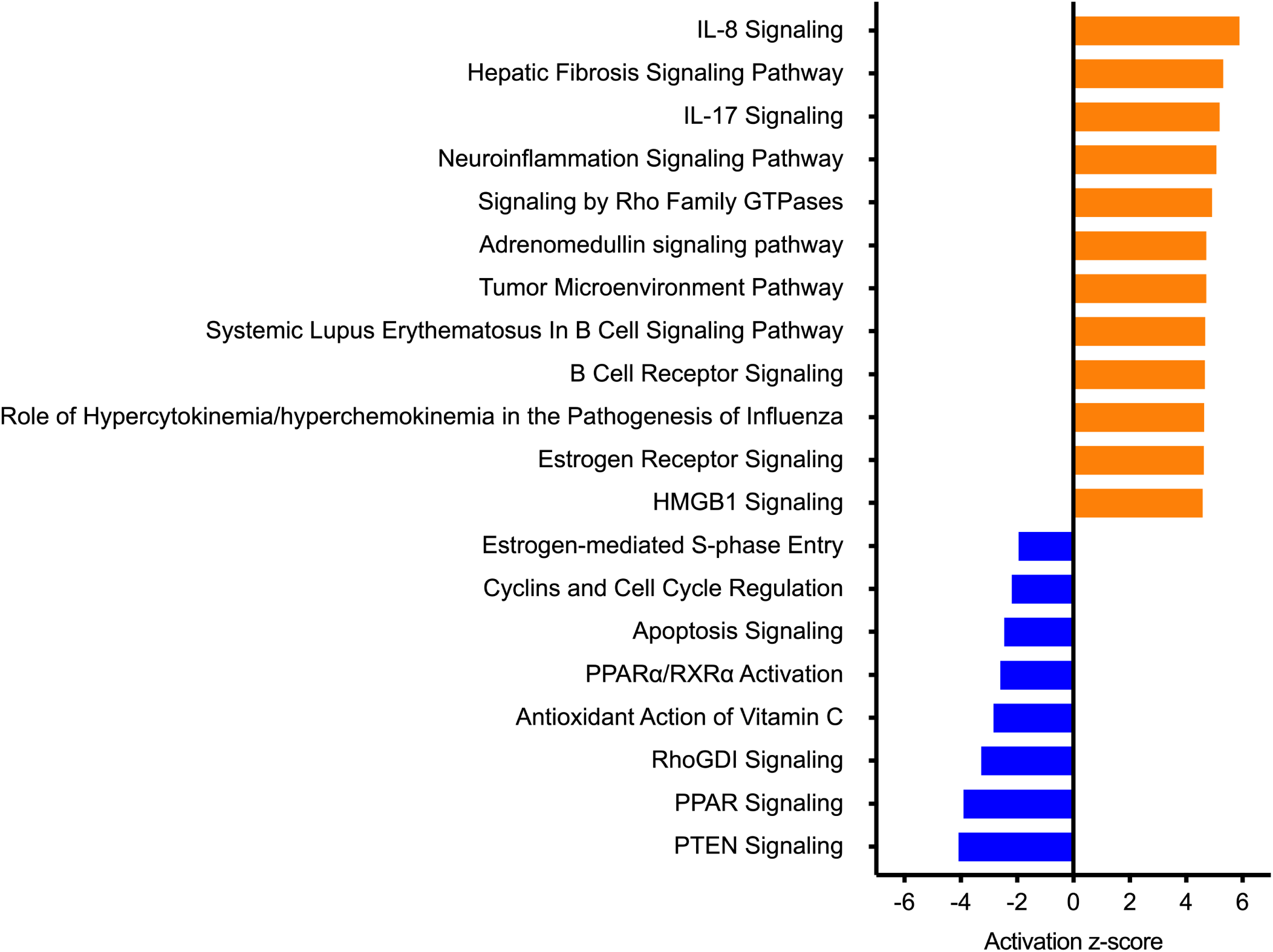
Host pathways potentially regulated by miRNAs during *C. burnetii* infection. Top 20 miRNA-targeted pathways significantly impacted in THP-1 macrophages infected with NMII. Orange bars show pathways that are putatively activated (z-score ≥ 1.5), and blue bars correspond to pathways that were predicted to be inhibited (z-score ≤1.5) using ingenuity pathway analysis (IPA). See full list in **Table S3**.

### miR-143-3p is down-regulated in *C. burnetii*-infected human alveolar macrophages

Among apoptosis-related miRNAs, we focused on miR-143-3p, which was significantly down-regulated in NMII-infected THP-1 cells (**Table 2, Figure 2A**). Because NMII is an avirulent laboratory strain and THP-1 cells are not natural host cells for *C. burnetii*, we measured miR-143-3p expression using primary human alveolar macrophages (hAMs) infected with the fully virulent *C. burnetii* Nine Mile RSA493 Phase I (NMI) strain. At 72 hpi, expression of miR-143-3p was significantly down-regulated in hAMs infected with *C. burnetii* (**Figure 2B**). Intriguingly, expression of miR-143-3p was significantly lower in NMI-infected than in NMII-infected hAMs. While the cause for this disparity is currently unknown, the full-length lipopolysaccharide (LPS) present in NMI might play a role because LPS is known to repress transcription of the miR-143/145 gene cluster (42).

**Figure 2.**
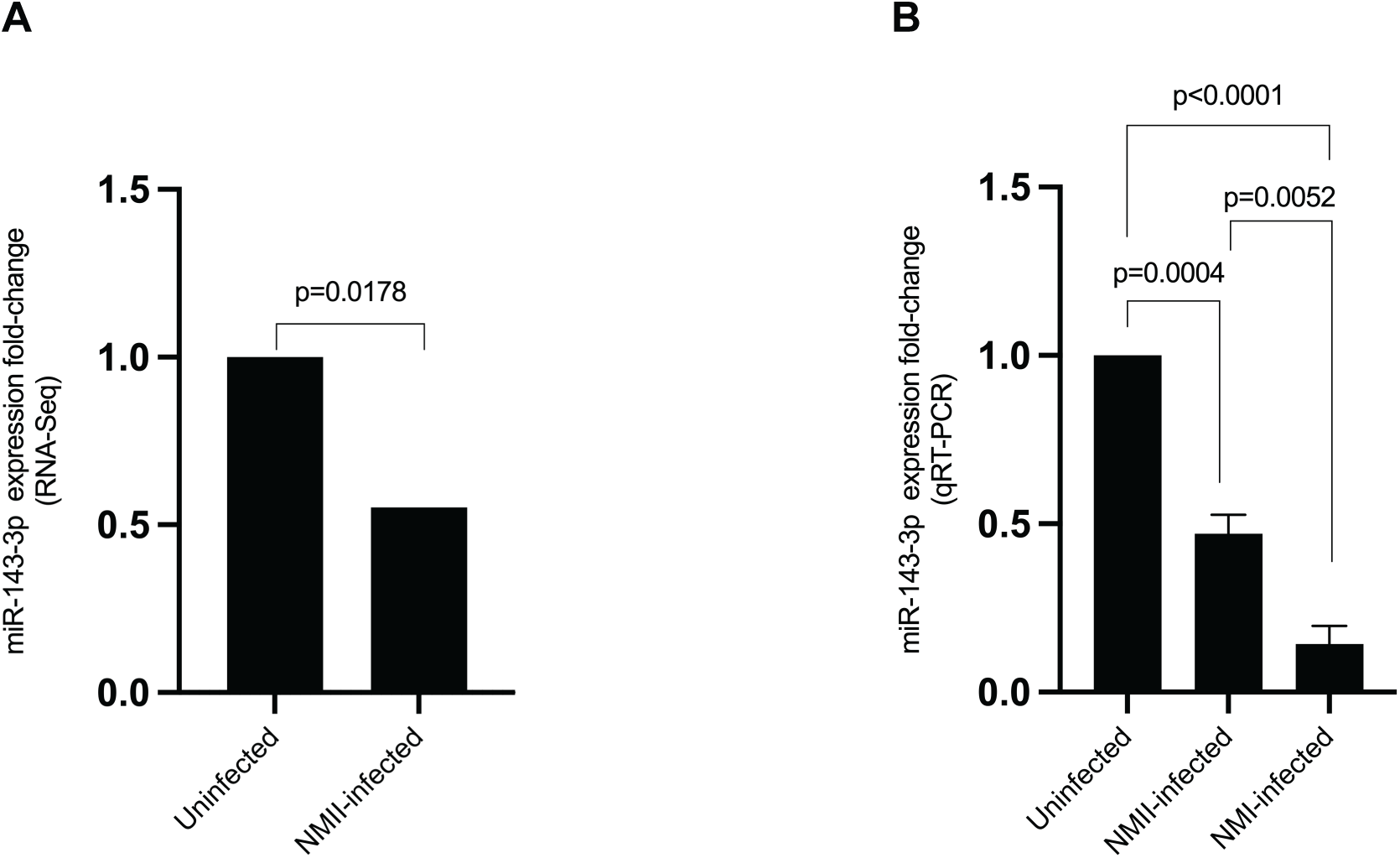
miR-143-3p expression is down-regulated in *C. burnetii*-infected macrophages. (**A**) Expression of miR-143-3p measured via RNA-seq (n = 3; 72 hpi) in NMII-infected THP-1 cells compared to uninfected cells. (**B**) Primary human alveolar macrophages (hAMs) infected with NMI or NMII or uninfected controls were analyzed for miR-143-3p expression using qRT-PCR at 72 hpi. Statistical significance in (A) was calculated using two-tailed paired Student’s t-test followed by Welch’s correction, and in (B) using one-way ANOVA followed by Tukey’s multiple comparison test (n = 3).

### Increased miR-143-3p expression inhibits *C. burnetii* growth

Because expression of miR-143-3p is down-regulated during *C. burnetii* infection, we tested how higher levels of miR-143-3p in host cells would impact intracellular growth. We first transfected HeLa cells with either miR-143-3p or a non-specific control miRNA (miR-control) and then infected cells with NMII and measured bacterial growth at 48 hpi. As shown in **Figure 3**, *C. burnetii* grew significantly better in untransfected and miR-control-transfected cells compared to miR-143-3p-transfected cells, indicating that an intracellular environment with low levels of miR-143-3p could be advantageous to the pathogen.

**Figure 3.**
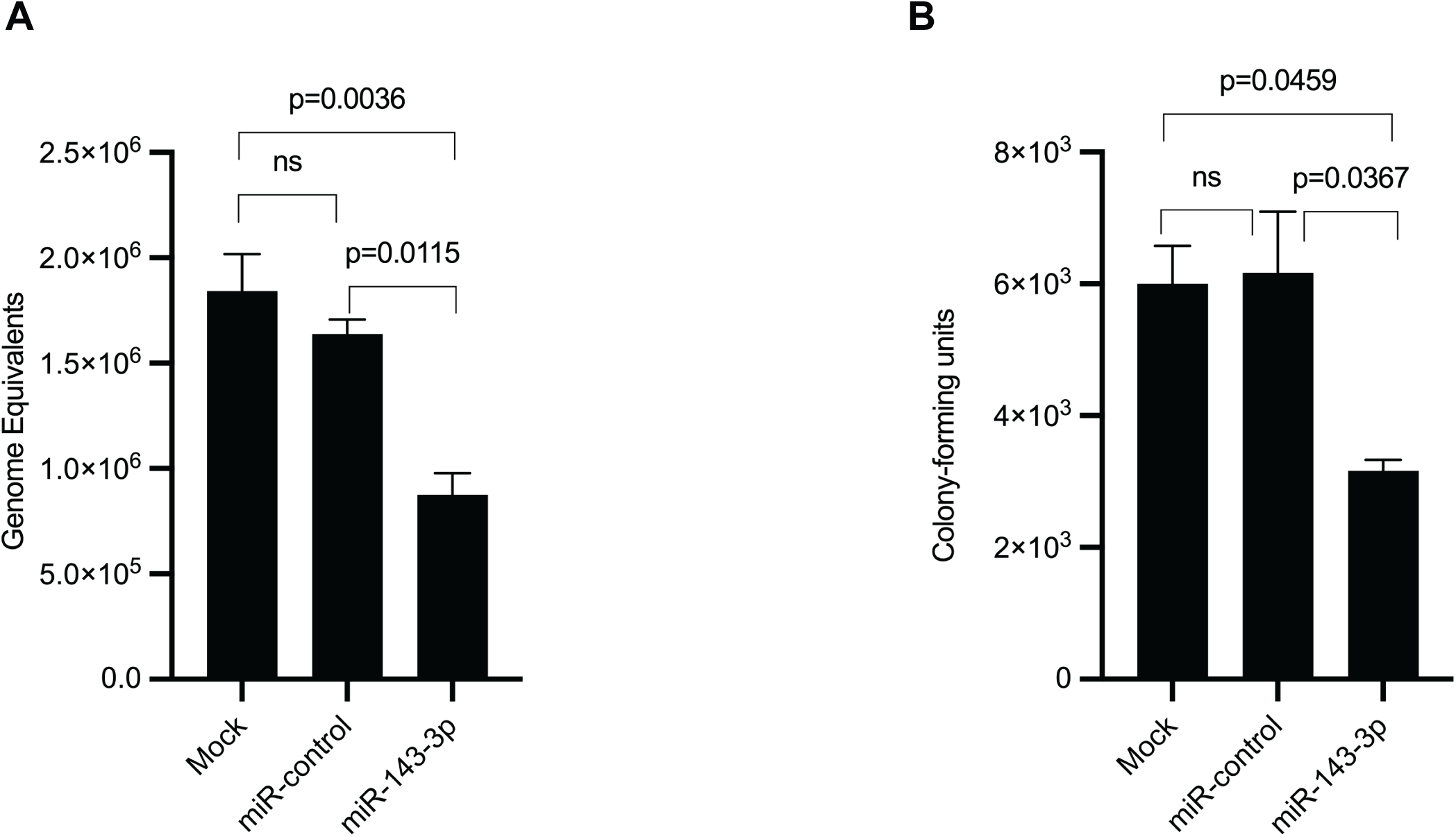
Intracellular growth of *C. burnetii* is inhibited in the presence of excessive miR-143-3p. Quantification of intracellular *C. burnetii* at 48 hpi using qPCR (**A**) or CFU assay (**B**) in miR-143-3p-transfected HeLa cells compared to untransfected cells (Mock) and cells transfected with non-specific control miRNA (miR-control). Statistical significance was determined using one-way ANOVA followed by Tukey’s multiple comparison test (ns: non-significant, n = 3).

### Early apoptosis is enhanced in cells transfected with miR-143-3p

To test the impact of miR-143-3p on apoptosis, we transfected HeLa cells with either miR-143-3p or miR-control and assessed early- and late-stage apoptosis using annexin V-PE and eFluor780 staining followed by flow cytometry (43). We observed that the percentage of early, but not late, apoptotic cells in the miR-143-3p-transfected population was significantly higher than in cells transfected with miR-control (**Figure 4**). To begin to understand miR-143-3p’s regulatory circuit, we assessed expression of *akt1* (AKT Serine/Threonine Kinase 1) and *bcl2* (B-cell lymphoma 2), two genes targeted by miR-143-3p that are central to apoptosis regulation in human macrophages (**Figure S1**) (44–47). Expression *of akt1* and *bcl2* genes and levels of activated Akt and Bcl-2 proteins were significantly reduced in miR-143-3p-expressing cells compared to cells transfected with miR-control (**Figure 5)**. Together, our data suggest that lower levels of miR-143-3p present in *C. burnetii*-infected cells increases expression of *akt1* and *bcl2*, which likely stalls apoptosis induction, thereby supporting the pathogen’s intracellular growth.

**Figure 4.**
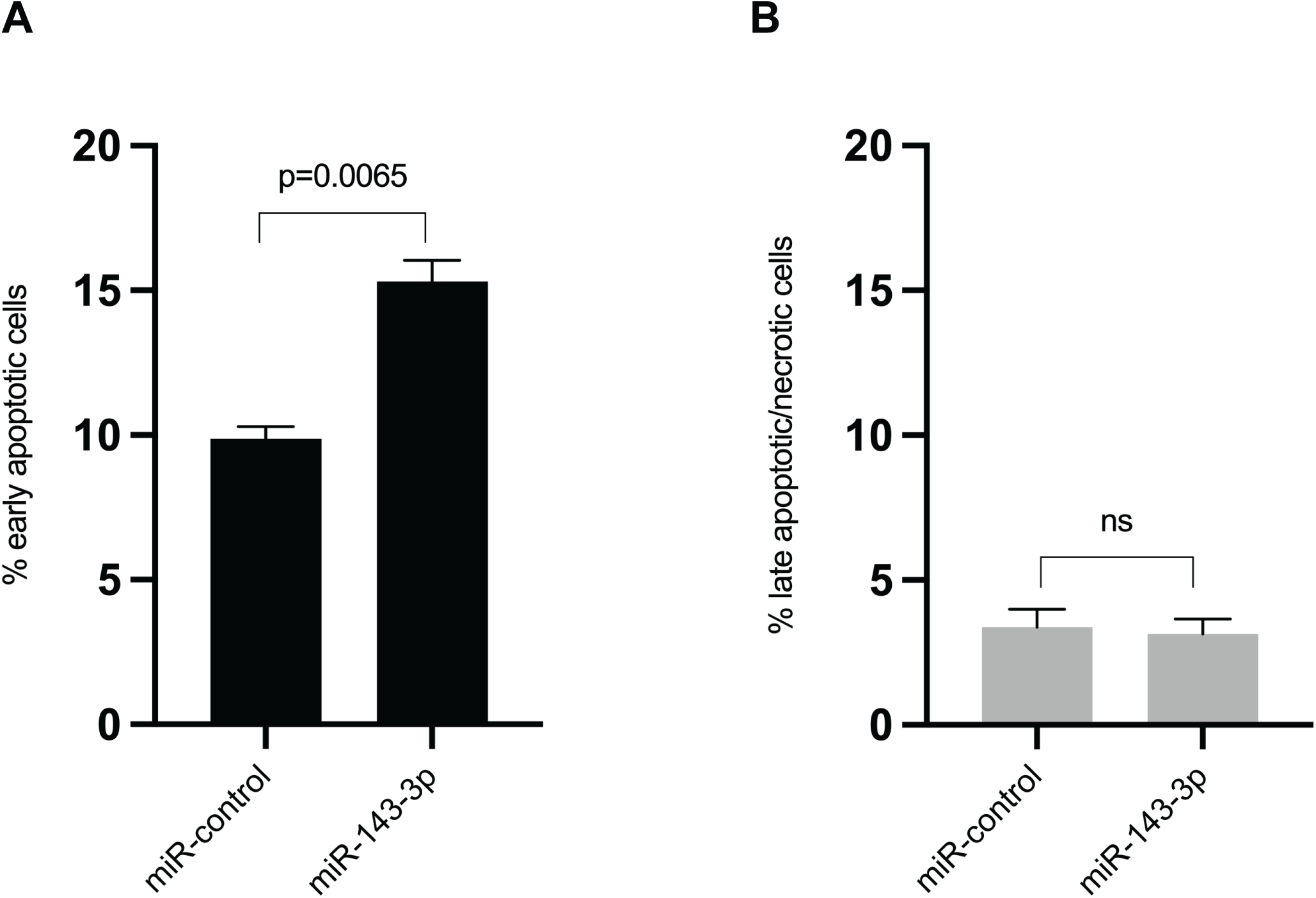
miR-143-3p transfection promotes early apoptosis. Percentage of early (**A**) or late (**B**) apoptotic HeLa cells transfected with either miR-143-3p or a non-specific control miRNA (miR-control). Early apoptosis (Annexin V-PE positive and eFluor780 negative) and late apoptosis/necrosis (Annexin V-PE positive and eFluor780 positive) were quantified by flow cytometry. Statistical significance was determined using two-tailed paired Student’s t-test followed by Welch’s correction (n = 3).

**Figure 5.**
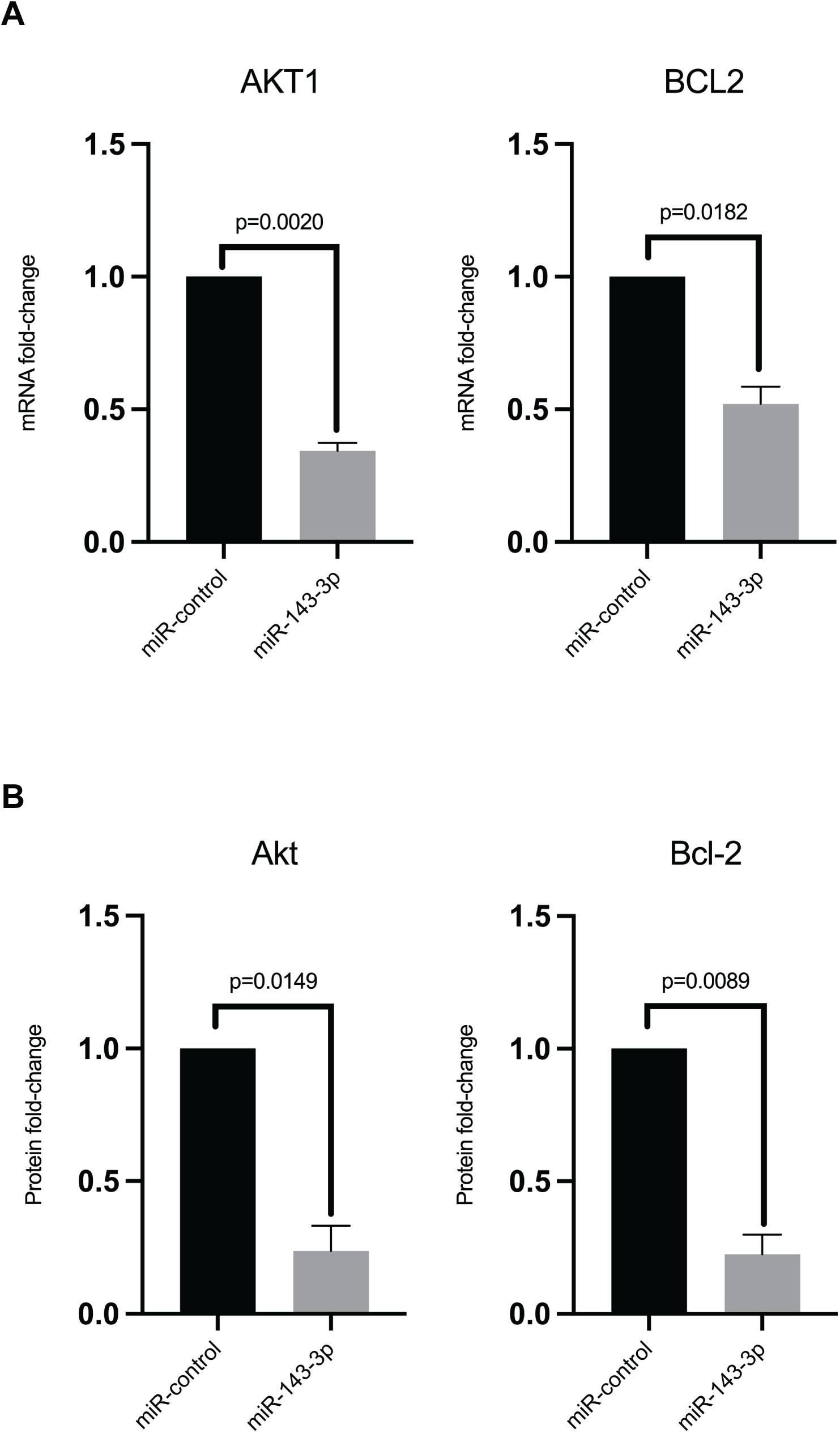
Increased miR-143-3p expression leads to reduced expression of apoptosis-related genes. (**A**) Expression (fold change) of *akt1* and *bcl2* genes in HeLa cells transfected with miR-143-3p compared to cells transfected with control miRNA (miR-control) measured using qRT-PCR. Statistical significance was determined using two-tailed paired t-test followed by Welch’s correction (n = 3). (**B**) Fold change in phosphorylated Akt (pSer473) and Bcl-2 (pSer70) proteins in HeLa cells transfected with miR-143-3p compared to cells transfected with miR-control measured using a multiplex immunoassay. Activated proteins were measured as median fluorescence intensity and statistical significance was determined using two-tailed paired Student’s t-test followed by Welch’s correction (n = 3).

### miR-143-3p has potential roles in autophagy

Analysis of miRNA-targeted pathways indicated that miRNAs could also be involved in autophagy **(Table S3)**, a process that is interconnected with apoptosis and is involved in the host response to *C. burnetii* (18, 48–50). We measured rapamycin-induced autophagic flux in HeLa cells transfected with either miR-143-3p or miR-control and observed that autophagic flux was slightly, but significantly, lower in miR-143-3p-transfected cells compared to miR-control-transfected cells (**Figure 6**). Several genes involved in autophagy, including *atp6v1a* (V-type proton ATPase catalytic subunit A) and *slc7a11* (Solute Carrier Family 7 Member 11) are controlled by miR-143-3p (24, 25). We observed significantly reduced expression of *atp6v1a* and *slc7a11* and concordant reduction in their encoded proteins (VATA and xCT, respectively) in miR-143-3p-transfected cells compared to control cells **(Figure 7, Table S4)**, indicating a role for these genes in reduced autophagic flux in miR-143-3p-expressing cells.

**Figure 6.**
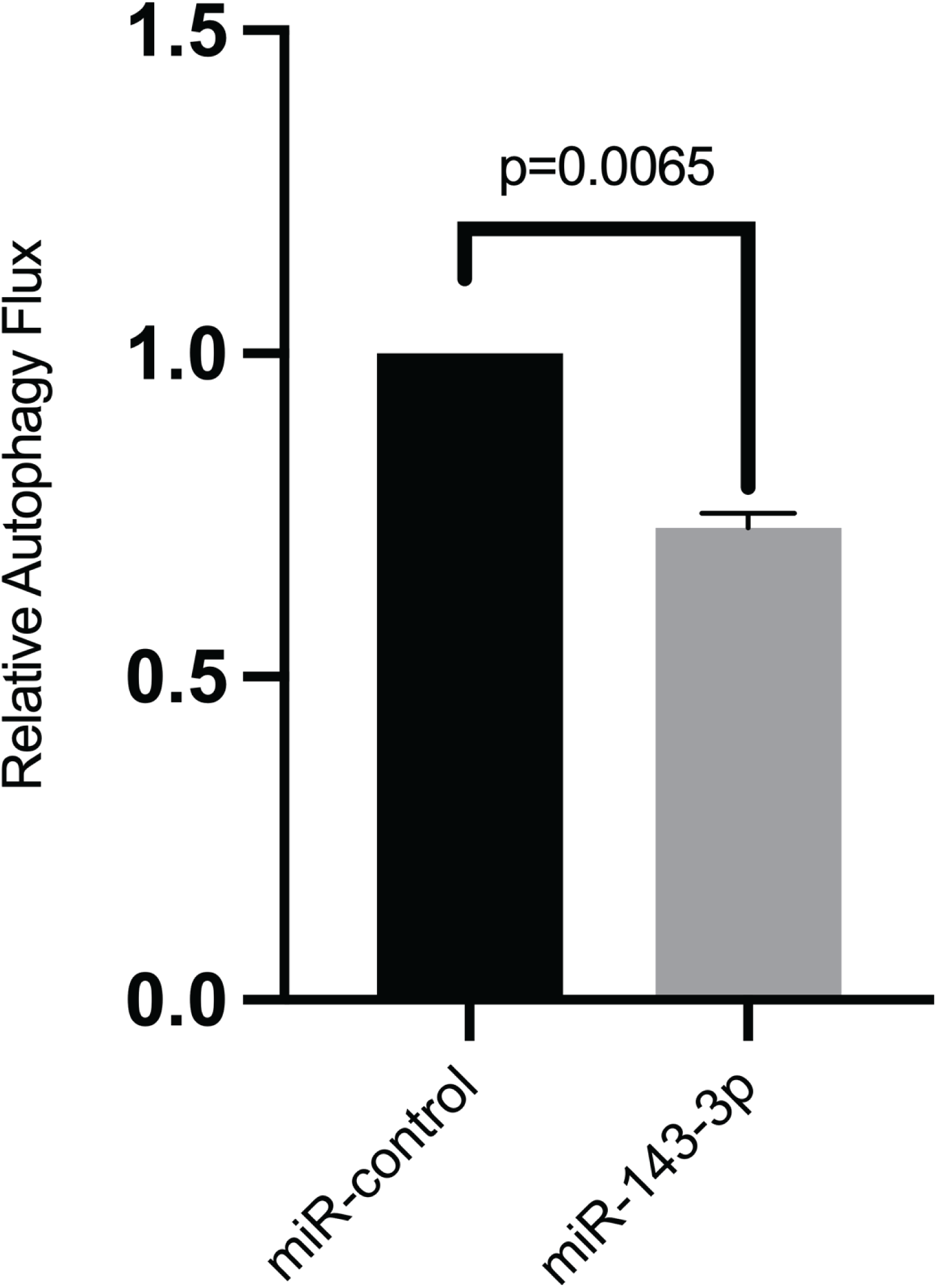
miR-143-3p inhibits autophagic flux. Y-axis shows relative autophagy flux reported as average brightness of CYTO-ID green (a cationic tracer that selectively labels autophagic compartments) per cell in miR-143-3p-transfected Hela cells compared to cells transfected with control miRNA (miR-control). Average CYTO-ID green brightness values were calculated from three sets of at least 200 cells per well and statistical significance was determined using two-tailed paired Student’s t-test followed by Welch’s correction (n = 3).

**Figure 7.**
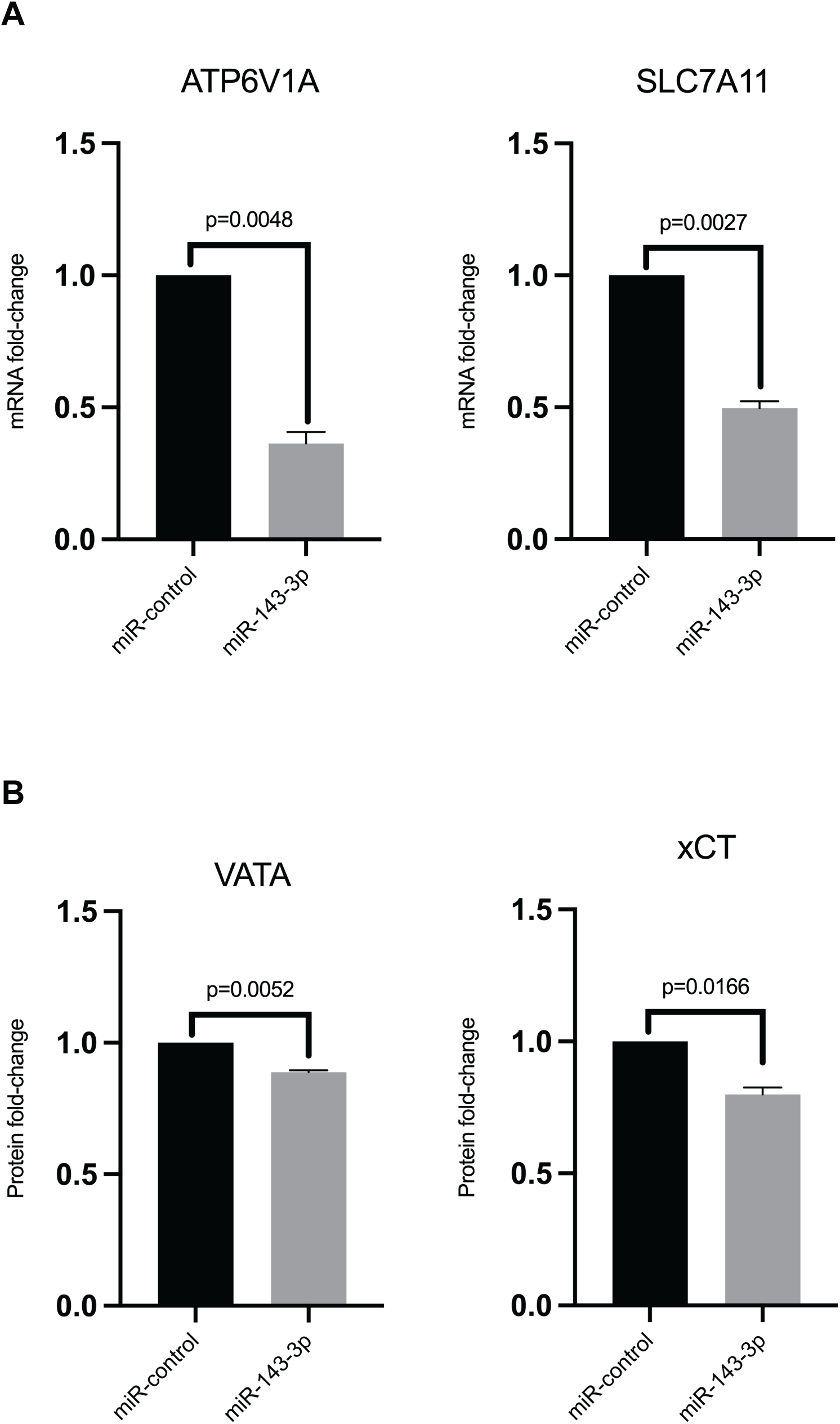
Transfection of HeLa cells with miR-143-3p decreased *atp6v1a*/VATA and *slc7a11*/xCT expression. **(A**) *atp6v1a* and *slc7a11* gene expression (fold change) in HeLa cells transfected with miR-143-3p compared to cells transfected with control miRNA (miR-control) measured using qRT-PCR. (**B**) VATA (encoded by *atp6v1a*) and xCT (encoded by *slc7a11*) proteins (fold change) in HeLa cells transfected with miR-143-3p compared to cells transfected with miR-control measured using quantitative mass-spectrometry. Statistical significance was determined using two-tailed paired Student’s t-test followed by Welch’s correction (n = 3).

## DISCUSSION

In this study, we show, for the first time, that *C. burnetii* infection elicits a robust miRNA-based host response, and closer examination of miR-143-3p function revealed that the miRNA promotes apoptosis and inhibits autophagy. This combination of phenotypes potentially produces an intracellular environment that is not conducive *to C. burnetii* growth, likely explaining down-regulation of miR-143-3p expression during *C. burnetii* infection. miR-143-3p presumably antagonizes *C. burnetii* growth by modulating components of the PI3K-Akt signaling network, which controls diverse host cell functions (**Figure S1**) (51). Activation of Akt by P13K induces expression of pro-survival Bcl-2 and inhibits pro-apoptotic proteins such as Bad (52), leading to decreased caspase-3 activation and subsequent inhibition of intrinsic apoptosis. During *C. burnetii* infection, Akt is activated, Bcl-2 is recruited to CCVs, and the pathogen secretes effector proteins that inhibit apoptosis (15, 16, 39, 53). Our results add to this knowledge by demonstrating that levels of Akt and Bcl-2 are significantly reduced and early apoptosis is significantly enhanced in cells transfected with miR-143-3p. No significant difference is evident in late apoptosis, a process characterized by DNA fragmentation, suggesting that miR-143-3p does not affect this process.

In addition to inhibiting apoptosis, *C. burnetii* manipulates autophagy to generate CCVs (10, 18, 49, 50). Interestingly, miR-143-3p appears to inhibit autophagy, and several genes involved in this process, including *atp6v1a* and *slc7a11*, are regulated by miR-143-3p (23–25, 54–57). Expression of both genes is significantly reduced in miR-143-3p-transfected cells, indicating that the miRNA’s effect on autophagy involves regulation of v-ATPase and cystine/glutamate antiporter xCT. In accordance with our results, inhibition of autophagy causes improper CCV maturation and reduces intracellular *C. burnetii* growth (17, 58, 59). Inhibition of ATP6V1A, a catalytic subunit of the lysosomal v-ATPase, likely disrupts autophagic flux by inhibiting V-ATPase-dependent acidification of developing CCV, thereby reducing *C. burnetii* growth (56). Furthermore, inhibition of xCT (encoded by *slc7a11*), a cystine/glutamate antiporter that relies on autophagy machinery to import cystine, could deprive the pathogen of cysteine, an amino acid essential to its growth (55, 60, 61). Finally, although this needs to be confirmed in macrophages, overexpression of miR-143-3p in human endothelial progenitor cells decreased LC3-II and increased p62 levels, two phenotypes that strongly imply a role for miR-143-3p in dampening autophagic flux (62). Cumulatively, our data suggest that down-regulation of miR-143-3p during *C. burnetii* infection promotes intracellular growth of the pathogen by delaying apoptosis and promoting autophagy. Further studies are required to determine if this is a *C. burnetii*-driven process and if LPS and/or T4SS effectors promote pathogen growth by down-regulating miR-143-3p expression.

## MATERIALS AND METHODS

### Bacterial strains and growth conditions

*Coxiella burnetii* Nine Mile RSA439 (Phase II, Clone 4) isolate (NMII) was cultured in acidified citrate cysteine medium-2 (ACCM-2) for 7 days at 37°C, 5% CO_2_, 2.5% O_2_ (63). Bacteria were quantified using PicoGreen (64, 65), collected by centrifugation (3000 x g,10 min, 4°c), resuspended in PBS containing 0.25 M sucrose (PBSS), and stored at -80°C until further use. Before infection, THP-1 cells (American Type Culture Collection, TIB-202) were differentiated into adherent, macrophage-like cells in RPMI-1640 supplemented with 1 mM sodium pyruvate, 0.05 mM beta-mercaptoethanol, 4500 mg/L glucose, and 10% heat-inactivated fetal bovine serum (FBS) at 37°C under 5% CO2 for 24 h using 30 nM phorbol 12-myristate 13-acetate (PMA), followed by 24 h of rest in PMA-free medium. Cells were infected with NMII at a multiplicity of infection (MOI) of 25 in serum-free growth medium for two hours and this time-point was considered to be 0 hpi. To remove extracellular bacteria, cells were washed three times with PBS followed by replacement with complete growth medium, which was replaced with fresh medium at 72 hpi. Primary human alveolar macrophages (hAMs) were harvested by bronchoalveolar lavage (BAL) from postmortem human lung donors and infected with either NMI (*C. burnetii* Nine Mile Phase I RSA493) or NMII at 25 MOI. hAMs were cultured at 37°C under 5% CO_2_ in Dulbecco’s modified Eagle/F-12 (DMEM/F12) medium (Gibco) containing 10% FBS for 72 hpi, as described previously (66).

### Transfection of HeLa cells

miRCURY LNA hsa-miR-143-3p and miRNA negative control were purchased from Qiagen. The negative control (miR-control) is a non-specific miRNA that shows no homology to any known miRNA or mRNA sequences annotated in mouse, rat, or human genomes. HeLa cells were reverse transfected with miRNAs as previously described (67). Briefly, 25 nM (final concentration) of either miR-143-3p mimic or miR-control was incubated in HiPerFect Transfection Reagent (Qiagen) for 10 min at room temperature to allow formation of transfection complexes. Transfection complexes were uniformly spotted at the bottom of each well of a 24-well tissue culture plate and 3.5 × 10^4^ HeLa cells were added to each well and incubated in OptiMEM medium. After 24 h, the medium was removed, and cells were washed twice with PBS before infection with NMII at an MOI of 100 in serum-free DMEM medium for two hours followed by washing and replenishing cells with serum-containing DMEM medium for experiments at 48 hpi.

### Intracellular growth assay

*C. burnetii* growth was measured at 48 hpi in HeLa cells transfected 24 h prior to infection with either miR-143-3p or miR-control. To quantify intracellular bacteria using qPCR, total DNA was extracted using QIAamp DNA Mini Kit (Qiagen) according to the manufacturer’s instructions and SYBR Green-based qPCR was performed using *C. burnetii*-specific primers (**Table S5**), as described previously (65). Quantification cycle (Cq) values were converted to bacterial genome equivalents (GE) using a standard curve, as described previously (64). Since qPCR-based quantitation does not differentiate between live and dead cells, we independently quantified viable intracellular bacteria by enumerating colony forming units (CFUs), as described previously (68, 69). Briefly, infected host cells were lysed in ice-cold water for 40 min at 4°C followed by repeated pipetting with a syringe and 25G needle to lyse remaining cells, centrifuged for 10 min at 70 x g (4°C) followed by centrifugation of supernatants for 1 min at 13,500 x g (4°C). Pellets were resuspended in ACCM-2, serially diluted, and spot-plated on ACCM-2 containing 0.5 mM tryptophan and 0.5% agarose. Plates were incubated for 10 days at 37°C, 5% CO_2_, and 2.5% O_2_ before enumerating CFUs.

### RNA sequencing, miRNA-target interactions and pathway analysis

THP-1 macrophages infected with NMII (MOI of 25) and uninfected controls were analyzed at 8, 24, 48, 72, and 120 hpi for miRNA and mRNA expression. At each time point, growth medium was replaced with 1 ml of TRI reagent (Life Technologies) and total RNA was extracted and treated with DNase (Invitrogen) per the manufacturer’s instructions. Samples were sequenced using Illumina NovaSeq 6000 for mRNAs and Illumina HiSeq 2500 for miRNAs at the Yale Center for Genome Analysis or Novogene Corporation, Sacramento, CA. Sequencing reads were mapped to the reference human miRNA (miRbase 22.1) or genome (GRCh38) databases using CLC Genomics Workbench v6.5 (Qiagen) (70, 71). Differential gene expression (log2 fold-change ≥ 0.75; adjusted p-value ≤ 0.05, Wald test) between NMII-infected and uninfected cells was calculated using DESeq2 (72). Inverse-expression pairings of differentially-expressed miRNAs and mRNAs and IPA core analysis of differentially-expressed genes were performed using Ingenuity Pathway Analysis (IPA, Qiagen) (36).

### Quantitative Reverse Transcription PCR (qRT-PCR)

Expression of 84 apoptosis-related miRNAs was measured using a miScript miRNA PCR Array Human Apoptosis kit (Qiagen). Briefly, total RNA extracted from infected or uninfected macrophages at 72 h pi (n = 3) were reverse transcribed using a miScript II RT kit and HiSpec buffer (Qiagen) and qPCR reactions were performed on a Stratagene Mx3005P Real-Time PCR system. miRNA expression data were normalized using the global quantitation cycle (Cq) mean of expressed miRNAs, and relative miRNA expression levels were calculated using the delta-delta Cq method (73). Expression of *akt1, bcl2, atp6v1a*, and *slc7a11* was quantified using qRT-PCR using *gapdh* primers as an endogenous reference control, as described previously (65). Briefly, total RNA was extracted, DNase-treated, and cDNA generated using RevertAid First Strand cDNA Synthesis Kit (Thermo Scientific). To perform qPCR, cDNA templates were diluted and mixed with gene-specific primers and SYBR Green (Applied Biosystems) in a 20 µl reaction according to the recommended protocol (Applied Biosystems). Primers used in this study are listed in **Table S5**. To assay miR-143-3p expression in hAMs, PCR primers for miR-143-3p and the endogenous reference *rnu*6 (small nuclear ribonucleic acid) were procured from Qiagen and qRT-PCR reactions were performed as described above.

### Apoptosis assay

Apoptosis was quantified in HeLa cells transfected with miR-143-3p or miR-control using Annexin V-PE (Invitrogen) and Fixable Viability Dye eFluor780™ (Invitrogen) (43). At 72 h post-transfection, cells were treated with trypsin, stained with efluor780 Fixable Viability Dye for 30 min in dark (4°C), followed by staining with Annexin V-PE for 15 min per the recommended protocol (Invitrogen). Cells were immediately assayed for early apoptosis (Annexin V-PE positive and eFluor780 negative) and late apoptosis/necrosis (Annexin V-PE positive and eFluor780 positive) markers using a FACSAria Fusion flow cytometer (BD Biosciences). Fluorescence parameters were gated using unstained and single-stained cells and a total of 10,000 events were counted for each sample using FACSDiva Software (BD Biosciences).

### Autophagic flux assay

CYTO-ID Autophagy Detection Kit 2.0 (ENZ-KIT175, Enzo Life Sciences) was used to quantify autophagic flux at 72 h post transfection on HeLa cells transfected with either miR-143-3p or miR-control. Cells were treated with media containing 200 nM rapamycin for 16 h to induce detectable levels of autophagy (74, 75), washed, and incubated for 30 min at 37°C in Microscopy Dual Detection Reagent containing CYTO-ID green detection reagent and Hoechst 33342 nuclear stain in 1X assay buffer. Three sets of at least 200 cells per well were immediately analyzed using a fluorescence microscope (Keyence Corporation) and levels of autophagic flux were measured as average CYTO-ID green brightness per cell calculated using Keyence BZ-X700 software (76, 77).

### Multiplex immunoassay

Activation (phosphorylation) of proteins involved in early apoptosis, including Akt (pS473) and Bcl-2 (Ser70), were quantified using 7-Plex Early Apoptosis Magnetic Bead Kit (EMD Millipore). Total cell lysate from HeLa cells transfected with miR-143-3p or miR-control was prepared using Lysis Buffer containing protease inhibitors, followed by bicinchoninic acid (BCA) quantitation at 72 h post-transfection. Briefly, 17.5 µg/well of diluted cell lysate was added to 1X magnetic beads at 1:1 ratio in a 96-well plate. The plate was incubated on a plate shaker (4°C, 700 rpm, dark) for 18 h, followed by washing and incubation with 1X Detection Antibody for 60 min at room temperature (RT) with shaking (700 rpm, dark). The detection antibody was then removed, and samples were incubated for 15 min at RT in the dark with 1X Streptavidin-PE (SAPE) followed by 15 min incubation (RT, dark) with the amplification buffer. SAPE and amplification buffer were removed, and beads were resuspended in 150 µl of assay buffer to analyze median fluorescence intensity (MFI) using a Luminex 200 system.

### Quantitative mass spectrometry

Total cell lysate from HeLa cells transfected with either miR-143-3p or miR-control were collected at 48h post-transfection in triplicates. TMT labeling and mass spectrometry were performed at the Proteomics Shared Resource facility at Oregon Health & Science University, as described previously (78). Briefly, samples were lysed, sonicated, and heated at 90°C for 10 min followed by overnight micro-digestion of each sample using an S-trap micro protocol. Peptides were labelled with TMT6-plex reagents, and multiplexed TMT-labeled samples were separated by two-dimensional reversed-phase-reversed-phase (2DRPRP) liquid chromatography on a Orbitrap Fusion Tribrid instrument (Thermo Scientific). Proteins were identified by searching against the human proteome in UniProt, and TMT reporter ion intensities were processed with in-house scripts. Differential protein abundance was determined by the Bioconductor package edgeR.

## Data availability

Sequencing reads from this study have been deposited on NCBI Sequence Read Archive (SRA) under the BioProject accession PRJNA679931.

## ACKNOWLEDGEMENTS

This work was supported in part by National Institutes of Health grants AI123464 and AI133023 to R.R.

## SUPPLEMENTARY FIGURE

**Figure S1. PI3K/Akt signaling network**. Activation of PI3K by pro-survival stimuli leads to phosphorylation/activation of Akt. Akt in turn activates anti-apoptotic proteins and pro-survival transcription factor NF-κB, which leads to the induction of pro-survival proteins Bcl-2 and Bcl-xL, and activation of XIAP. Bcl-2 prevents the release of cytochrome c from mitochondria, thereby preventing apoptosis. Inhibition of *akt1* and *bcl2* expression by miR-143-3p (44–47), shown in red, could reverse this process to promote apoptosis.

## SUPPLEMENTARY TABLES

**Table S1**. Differentially expressed miRNAs and mRNAs in NMII-infected THP-1 cells.

**Table S2**. Inverse expression pairs of differentially expressed miRNAs and target mRNAs.

**Table S3**. List of pathways enriched for miRNA-regulated genes.

**Table S4**. List of downregulated proteins in miR-143-3p-transfected HeLa cells identified using mass spectrometry.

**Table S5**. List of primers used in this study.

